# Dissecting β-Cardiac Myosin and Cardiac Myosin-Binding Protein C Interactions using a Nanosurf Assay

**DOI:** 10.1101/2022.03.11.483820

**Authors:** Anja M. Touma, Wanjian Tang, David V. Rasicci, Duha Vang, Ashim Rai, Samantha B. Previs, David M. Warshaw, Christopher M. Yengo, Sivaraj Sivaramakrishnan

**Affiliations:** Department of Genetics, Cell Biology, and Development, University of Minnesota, Minneapolis, MN, USA; Department of Cellular and Molecular Physiology, Penn State College of Medicine, Hershey, Pennsylvania, USA; Department of Molecular Physiology and Biophysics, Cardiovascular Research Institute, University of Vermont, Burlington, VT

## Abstract

Cardiac myosin-binding protein C (cMyBP-C) regulates cardiac contractility by slowing shortening velocity and sensitizing the thin filament to calcium. cMyBP-C has been shown to interact with the proximal myosin S2 tail and the thin filament. However, the relative contribution of these interactions to the collective modulation of actomyosin ensemble function remains unclear. Hence, we developed a “nanosurf” assay as a model system to interrogate cMyBP-C interactions with actin and/or myosin. Synthetic thick filaments were generated using recombinant human β-cardiac myosin subfragments (HMM or S1) attached to DNA nanotubes, with 14 or 28 nm spacing, corresponding to the 14.3 nm myosin spacing found in native thick filaments. *In vitro* motility assays with myosin bound to the surrounding surface, exhibit enhanced thin filament interactions with synthetic thick filaments. No significant differences were observed in mean thin filament velocities between 14 and 28 nm spacing, consistent with our previous results for myosin V, VI, and β-cardiac myosin S1. Our nanosurf assay demonstrates the slowing of actomyosin motility by cMyBP-C. Alternating β-cardiac myosin HMM and cMyBP-C N-terminal fragments, C0-C2 or C1-C2, every 14 nm on the nanotube, reduced the mean thin filament velocity 4-6 fold relative to myosin alone. Interestingly, similar inhibition was observed using a β-cardiac myosin S1 construct, which lacks the S2 region proposed to interact with cMyBP-C, suggesting the actin-cMyBP-C interactions may dominate the inhibitory mechanism. No significant inhibition of thin filament velocity was observed with a C0-C1f fragment, lacking the majority of the M-domain, supporting the importance of this domain for inhibitory interaction(s). A phosphomimetic C0-C2 fragment showed a 3-fold higher velocity compared to its phosphonull counterpart, further highlighting phosphorylation-dependent regulation via the M-domain. Together, we have established the nanosurf assay as a tool to precisely manipulate spatially dependent cMyBP-C binding partner interactions, shedding light on the molecular regulation of β-cardiac myosin contractility.

**STATEMENT OF SIGNIFICANCE:** Cardiac myosin-binding protein C (cMyBP-C) is the most frequently mutated protein associated with hypertrophic cardiomyopathy (HCM), a common cause of sudden cardiac death. Despite the importance of cMyBP-C in cardiac contractility, the mechanisms underlying this regulation are unclear due to experimental challenges in studying the complex, transient, weak interactions of cMyBP-C with the contractile proteins of the sarcomere. In this study, we created a nanosurf synthetic DNA thick filament assay to dissect the cMyBP-C interactions with actin and human β-cardiac myosin. We demonstrate actomyosin inhibition by cMyBP-C fragments regardless of recombinant human β-cardiac myosin subfragment (HMM or S1) and highlight the importance of the cMyBP-C M-domain using cMyBP-C fragments and phosphomimetics.

## INTRODUCTION

Cardiac myosin-binding protein C (cMyBP-C) is a large, multidomain thick filament-associated sarcomeric protein that modulates cardiac muscle contractility by tuning the speed and efficiency of contraction and relaxation (1–3). Despite almost 50 years since its discovery (4, 5), the mechanisms by which cMyBP-C modulates contractility, including its binding partners and the associated regulatory mechanisms, are not completely understood. The importance of cMyBP-C in regulating contraction was first appreciated in the 1990s when two mutations in *MYBPC3*, the gene encoding cMyBP-C, were shown to cause hypertrophic cardiomyopathy (HCM), a heritable heart condition characterized by left ventricular hypertrophy and diastolic dysfunction (6, 7). Since then, over 350 *MYBPC3* mutations have been identified in HCM patients, and it has become the most prominent gene linked to HCM (8). The discovery of these mutations-including nonsense, frameshift, and missense mutations distributed across the protein-have provided substantial motivation towards understanding the role of cMyBP-C in the sarcomere (3). However, despite this increased interest and established importance of cMyBP-C in regulating contraction, the mechanisms by which cMyBP-C regulates contraction have largely remained elusive due to experimental challenges in studying the complex, transient, weak interactions of cMyBP-C in the context of the sarcomere.

Existing studies suggest that cMyBP-C modulates cardiac muscle contractility by sensitizing the thin filament to calcium, while slowing shortening velocity through interactions with actin and/or myosin (9–11). cMyBP-C exists in two regions of the muscle sarcomere A-band, in 7-9 stripes spaced 43 nm apart. This aligns with the 43 nm helical myosin head repeat on the thick filament (12–14). cMyBP-C contains 11 subdomains, named sequentially C0-C10, and is anchored to the thick filament backbone at its C-terminus by a high-affinity LMM myosin binding site in C10 (Supplemental Figure 1a) (15, 16). The N-terminus extends away from the thick filament and engages in regulatory interactions with myosin and/or actin (10, 17, 18). The cardiac isoform contains a regulatory cMyBP-C motif or M-domain between C1 and C2 domains, which co-sedimentation assays have established to be the primary interacting region for myosin S2 (19–23). Studies with actin have found that the M-domain, as well as the C0, C1, and C2 domains, interact with actin (21, 24–27). The N- and C-terminal cMyBP-C interactions to the thin and thick filaments, respectively, may tether the two filaments thus functioning as a brake on shortening velocity at high calcium concentrations (1, 17). Despite this wealth of information, the relative significance of actin vs myosin interactions with distinct N-terminal cMyBP-C domains is still a matter of debate. Additionally, cMyBP-C function itself may be regulated, as the M-domain contains four serines that are thought to be phosphorylated hierarchically to disrupt cMyBP-C interactions with either actin filaments (28–30) or myosin (31, 32). Phosphorylation of the M-domain is considered a regulatory mechanism in response to β-adrenergic stimulation (32).

The complex interactions of cMyBP-C with actin and myosin have made *in vitro* assessment of cMyBP-C challenging. Some *in vitro* motility studies have demonstrated inhibition of actomyosin motility by a variety of cMyBP-C N-terminal fragments (30, 33). However, mechanistic interpretation of these assays is limited by a lack of control over protein stoichiometry and orientation. Likewise, single molecule and solution assays do not capture the unique architecture of motor ensembles or cMyBP-C interactions in the context of the sarcomere. To overcome these limitations, we utilized DNA nanotechnology to build a synthetic thick filament that would allow systematic *in vitro* dissection of cMyBP-C interactions, with control over protein type, stoichiometry, and spacing (Figure 1a; Supplemental Figure 1) (34). We incorporated these synthetic thick filaments in a standard *in vitro* motility assay to promote thin filament interactions with the decorated DNA nanotubes. Using this nanosurf assay, we characterized the cMyBP-C domains and interactions that modulate acto-myosin motility.

**Figure 1:**
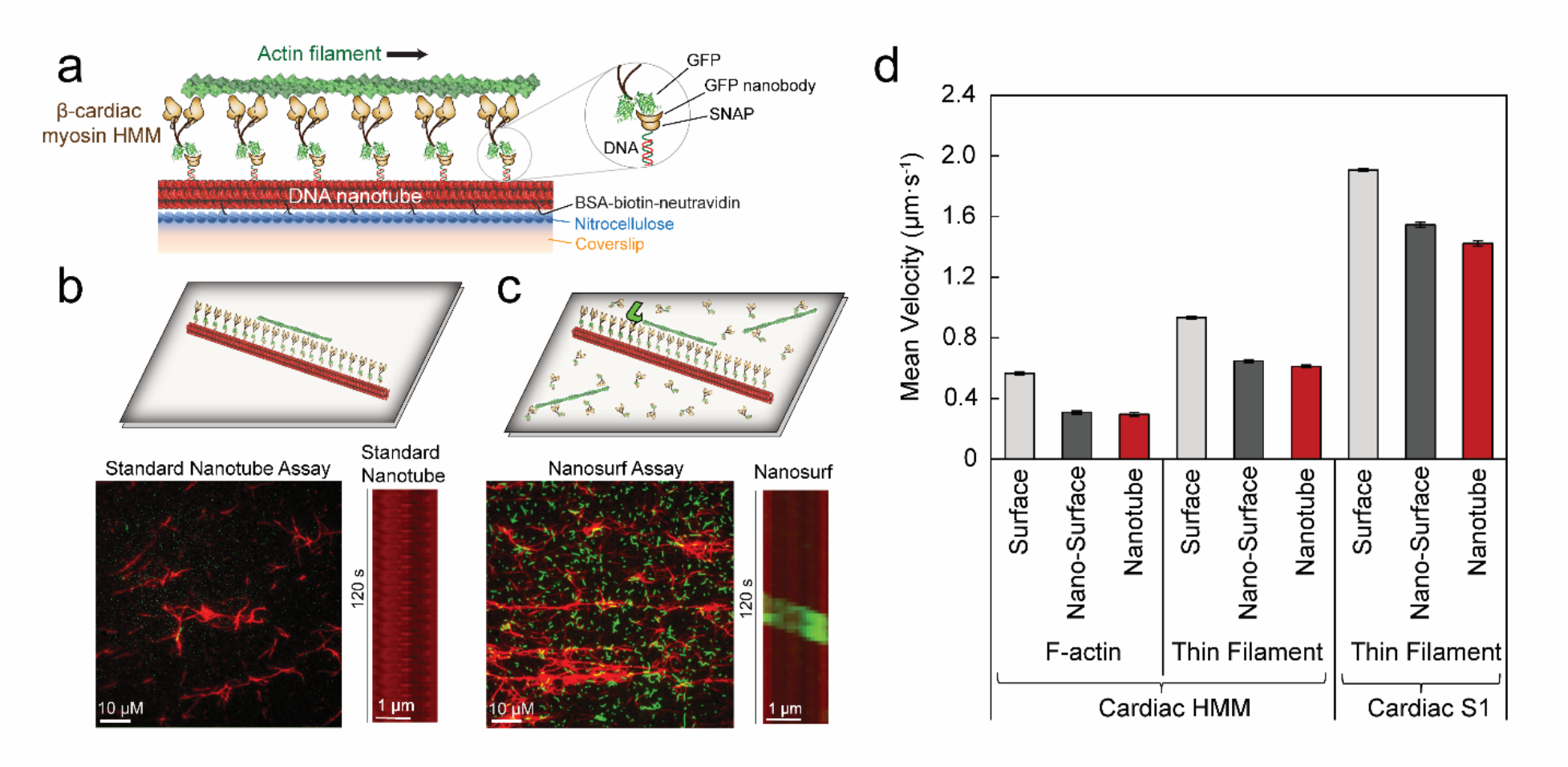
Optimization of Human β-Cardiac Myosin HMM Nanosurf Assay. **a**, Schematic of the β-cardiac myosin synthetic thick filament assay with recombinant human β-Cardiac myosin HMM (*brown*) with C-terminal GFP attached to the nanotube (*red*) via GFP nanobody-SNAP labeled with complementary oligo to the nanotube DNA handle (*inset*). Nanotubes are attached to the nitrocellulose-coated (*blue*) coverslip (*orange*) via BSA-biotin-neutravidin (*black X*’*s*) **b**, Schematic of standard nanotube assay within a flow cell with motor bound to nanotubes only and nanotubes attached to a coverslip surface blocked with BSA (*top*). Field of view at 1000x (*bottom left*) and kymograph (*bottom right*) for standard nanotube assay (*nanotubes in red)* with β-cardiac myosin HMM showing the low landing rate of actin filaments *(green)* on the nanotube surface. **c**, Schematic (*top*) of nanosurf assay with motor bound to both the nanotubes and the coverslip surface and actin (*green*) can be observed gliding from the coverslip surface onto the nanotube (*green arrow*). Field of view at 1000x (*bottom left*) and kymograph (*bottom right*) for nanosurf assay with β-cardiac myosin HMM showing a higher landing rate for actin filaments on the nanotube surface. **d**, Velocities of F-actin (*left*) and regulated thin filaments (*middle*) with β-cardiac myosin HMM and regulated thin filaments with β-cardiac myosin S1 (*right*) on a standard *in vitro* motility surface (*light grey*), on the coverslip surface in the nanosurf assay (nano-surface; *dark grey*), and on the nanotubes in the nanosurf assay (*red*). Mean velocities represented as μm·s^−1^ ± SE. N = 132-390 filaments from 3-5 independent protein preparations per condition.

## METHODS

### DNA Nanotube and Benzyl-Guanine Oligonucleotide Preparation

Cy5-labeled ten-helix DNA nanotubes composed of 40 single-stranded DNA tiles with 14 or 28 nm spacing between single-stranded protein binding handles were prepared with biotin strands for surface attachment using an annealing protocol as previously described (34). The nanotube DNA handles at 14 or 28 nm are designed to anneal to DNA strands bound to a SNAP protein encoded with the GFP nanobody. The SNAP protein binds benzyl-guanine treated DNA oligos. To prepare the benzyl-guanine oligo, benzyl-guanine NHS ester (BG-GLA-NHS; New England Biolabs, Ipswich, MA) was covalently linked to C6-amine oligonucleotides (with or without Cy3 modification, C6-amine-Cy3-a’ or C6-amine-a’). This was accomplished by incubating 0.17 mM C6-amine-a’ or C6-amine-b’ with 11.6 mM BG-GLA-NHS in 0.1 M sodium borate, pH 8.5 at 37°C for ~4 hours with rotation. The BG-labeled oligonucleotide was then purified twice through Illustra G-50 micro columns (GE Healthcare, Buckinghamshire, UK) equilibrated in 2 mM Tris, pH 8.5 since the primary amine reacts with any unreacted benzyl guanine. The final BG-labeled oligonucleotide concentration was determined by measuring Cy3 absorbance (for BG-Cy3-a’ or BG-Cy3-b’) or by estimating the concentration of single-stranded DNA (for BG-a’ or BG-b’) using a NanoDrop spectrophotometer (NanoDrop One^C^, ThermoFisher, Waltham, MA).

### GFP Nanobody–SNAP and Cardiac Myosin Binding-Protein C (cMyBP-C)-SNAP Preparation

The DNA sequence for the GFP-nanobody was generated and synthesized by Genewiz (South Plainfield, NJ) as previously described (35). The SNAP-tagged GFP nanobody construct contained from N- to C-terminus: the GFP nanobody, a flexible (Gly-Ser-Gly)2 linker, the SNAP tag for oligonucleotide labeling, and both FLAG and 6xHis purification tags. N-terminal fragments (C0-C2, C0-C1f, or C1-C2) of the *Mus musculus* cardiac myosin-binding protein C cDNA were cloned into pBiex-1 and engineered with a C-terminal SNAP-FLAG-6xHis. C0-C2 and C1-C2 phospho-null or phospho-mimetic constructs were created by mutating S273, S282, S302, and S307 to alanines or aspartic acids, respectively. For cMyBP-C constructs containing a linker, either a 10 nm ER/K α-helical linker (derived from *Sus scrofa’s* Myosin VI) or a 30 nm linker (derived from the Kelch-motif family protein, *Trichomonas vaginalis*), as previously characterized and described, were inserted between the cMyBP-C C-terminus and the N-terminus of the SNAP (36–38). Gly-Ser-Gly repeats were inserted between each element to ensure rotational flexibility. The C0-C2 and C1-C2 fragments contained all structural elements within and including their respective domains, notably including the M-domain between C1 and C2. The C0-C1f fragment contained the C0 and C1 domains with the respective unstructured pro-ala rich linker region, as well as the first 17 amino acids of the M-domain, as described previously (30). The proteins were then transiently transfected (pBiex-1; Escort IV, Sigma-Aldrich, St. Louis, MO) into Sf9 cells, expressed, and affinity purified similarly to previously published protocols (35, 39). Briefly, cells were lysed at 72 h in lysis buffer: 0.5% IGEPAL, 4 mM MgCl_2_, 200 mM NaCl, 7% sucrose, 20 mM Imidazole (pH 7.5), 0.5 mM EDTA, 1 mM EGTA, 1 mM dithiothreitol (DTT), 1 μg/ml phenylmethylsulfonyl fluoride (PMSF), 10 μg/ml aprotinin, and 10 μg/ml leupeptin. Lysates were centrifuged (176,000*g*, 4°C, 25 min) in a TLA 100.4 rotor (Beckman Coulter, Brea, CA) and bound to anti-FLAG M2 affinity resin (Sigma-Aldrich, St. Louis, MO) at 4°C. The FLAG resin was washed with wash buffer: 20 mM imidazole, 150 mM KCl, 5 mM MgCl_2_ ,1 mM EDTA, 1 mM EGTA, 1 mM DTT, 1 μg/mL PMSF, 10 μg/mL aprotinin, and 10 μg/mL leupeptin, pH 7.4. To label the SNAP tag, GFP nanobody or cMyBP-C fragments bound to anti-FLAG resin were incubated with excess (>10 μM) BG-oligonucleotide in wash buffer overnight at 4°C with rotation. BG-oligonucleotide-labeled SNAP proteins bound to resin were again washed with wash buffer and eluted using 0.2 mg/mL FLAG peptide (Sigma-Aldrich, St. Louis, MO) overnight at 4°C with rotation. To confirm labeling, proteins were loaded onto a 10% SDS gel followed by Coomassie staining. The protein was stored in 50% glycerol (v/v) at −20°C.

### Recombinant Human β-cardiac Myosin Preparation

The human β-cardiac myosin cDNA (AAA51837.1) was purchased from Thermo Fisher Scientific, Waltham, MA. PCR amplification was used to subclone the M2β-HMM fragment containing the first 25-heptads of the proximal S2 region (1-1016) into a pshuttle vector (a gift from Dr. Don Winkelmann). A GCN4 leucine zipper was added after the S2 region to promote dimerization which was followed by an eGFP tag at the C-terminus. An N-terminal FLAG tag was also included for purification. Similarly, an M2β-S1 (1-842) construct with a C-terminal eGFP tag and N-terminal FLAG tag was generated and subcloned into a shuttle vector provided by Vector Laboratories, Burlingame, CA. Recombinant adenovirus was generated as previously described to express M2β-HMM-eGFP in C_2_C_12_ cells (40). Vector Laboratories produced the initial recombinant adenovirus of M2β-S1-eGFP. High-titer adenovirus was produced by a method developed in the Winkelmann laboratory (41, 42) and as previously described (40, 43). For expression of β-cardiac myosin eGFP, C_2_C_12_ cells (typically 20-30, 145 mm diameter plates) were differentiated and infected with recombinant adenovirus (5 x 10^8^ PFU/ml) diluted into differentiation media as described previously (40). The cells were harvested on day 10 and myosin eGFP was purified by FLAG affinity chromatography as described. The eluted M2β-HMM-eGFP or M2β-S1-eGFP was then ammonium sulfate precipitated and dialyzed into MOPS20 buffer overnight at 4°C. M2β-HMM-eGFP and M2β-S1-eGFP was assessed for purity by coomassie stained SDS-polyacrylamide gels and concentrations were determined by eGFP absorbance (_*ε*488_ = 55,000 M^−1^×cm^−1^) or Bradford assay using BSA as a standard.

### Actin and Reconstituted Regulated Thin Filament Preparation

For assays with F-actin, actin was purified from rabbit skeletal muscle using an acetone powder method (44). The actin concentration was determined by absorbance at 290 nm (ε290= 2.66 x 104 M-1·cm-1). A molar equivalent of tetramethylrhodamine (TRITC) phalloidin (MilliporeSigma, Burlington, MA) was added to stabilize and fluorescently label the F-actin. Calcium-regulated native thin filaments were prepared from mouse ventricular tissue and TRITC-labeled as described previously (9).

### Standard *In Vitro* Motility Surface Assay

Flow chambers were created using tape placed ~3 mm apart on a glass slide to adhere coverslips coated with 0.1% collodion in amyl acetate (Electron Microscopy Sciences, Hatfield, PA). For standard gliding assays with β-cardiac myosin eGFP, myosin was attached to coated coverslips through GFP nanobody with an encoded SNAP protein bound to oligonucleotides annealed to single-stranded DNA nanotube handles (35). Briefly, GFP nanobody SNAP was incubated in flow chambers at ~0.5-1 μM for 4 minutes. Excess GFP nanobody SNAP was washed out, and the surface was incubated with assay buffer (AB: 20 mM Imidazole, pH 7.5, 25 mM KCl, 4 mM MgCl2, 1 mM EGTA, 1 mM DTT) and BSA (AB.BSA: AB + 1 mg/mL BSA) for 2 minutes. Finally, human β cardiac myosin at ~0.6-0.8 μM was incubated on the surface for 4 minutes and washed out with AB.BSA (for experiments with regulated thin filaments, flow cell was washed with pCa 5 buffer containing an appropriate CaCl2 concentration calculated using Max Chelator from UC-Davis). The final actin imaging solution was added containing tetramethylrhodamine (TRITC) phalloidin-labeled (Invitrogen, Waltham, MA) F-actin (or regulated thin filaments in pCa 5 buffer), 0.1% methylcellulose, 2 mM ATP, 1 mM phosphocreatine, 0.1 mg/mL creatine-phosphokinase, 45 μg/mL catalase, 25 μg/mL glucose oxidase, and 1% glucose. Flow cells were imaged at 1000x magnification on a Nikon (Tokyo, Japan) TiE microscope equipped with a 100x 1.4 NA Plan-Apo oil-immersion objective, 1.5x magnifier, Nikon Perfect Focus System, mercury arc lamp, Evolve EMCCD camera (512 pixel x 512 pixel; Photometrics), and Nikon NIS-Elements software. All *in-vitro* motility assays were performed at room temperature (20-23°C).

### Standard Nanotube Assay

The standard nanotube assay, previously reported and characterized (34), was presented for comparison in Figure 1b. At room temperature (20-23°C), nanotubes were attached to the coverslip surface coated with 0.1% collodion in amyl acetate (Electron Microscopy Sciences, Hatfield, PA) using biotinylated-BSA at 0.1 mg/mL in AB (AB: 20 mM Imidazole, pH 7.5, 25 mM KCl, 4 mM MgCl2, 1 mM EGTA, 1 mM DTT) incubated for 4 minutes. Excess biotin-BSA was washed out and the surface was incubated with AB.BSA (AB + 1 mg/mL BSA) for 2 minutes. Next, neutravidin at 0.1 mg/mL in AB.BSA was incubated for 4 minutes. AB.BSA was added to wash out excess neutravidin. Nanotubes were added at 2-5 nM concentration in AB.BSA.nt (AB.BSA + 5–10 nM random DNA nucleotide mix to reduce non-specific interactions) and incubated for 4 minutes. Excess nanotubes were washed out of the chamber with AB.BSA.nt. For attachment of β-cardiac myosin with eGFP, GFP nanobody SNAP at ~0.5-1 μM in AB.BSA.nt was incubated in flow chambers for 4 minutes. Excess GFP nanobody SNAP was washed out, and the surface was incubated with AB.BSA.nt for 2 minutes. Human β-cardiac myosin at ~0.6-0.8 μM was incubated on the surface for 4 minutes and washed out with AB.BSA.nt. The final actin imaging solution was added containing sheared tetramethylrhodamine (TRITC) phalloidin-labeled (Invitrogen, Waltham, MA) F-actin (sheared to ensure actin filament lengths are shorter than ~5 μm nanotubes), 0.1% methylcellulose, 2 mM ATP, 1 mM phosphocreatine, 0.1 mg/mL creatine-phosphokinase, 45 μg/mL catalase, 25 μg/mL glucose oxidase, and 1% glucose. For each flow cell, the actin channel was imaged as above at 0.5-3 second intervals for 2-3 minutes. For each actin video, several nanotube images were acquired of the same field of view for overlay.

### Nanosurf Assay

Flow chambers were created with nitrocellulose-coated coverslips as described above. At room temperature (20-23°C), nanotubes were attached to the coverslip using biotinylated-BSA at 0.1 mg/mL in AB (AB: 20 mM Imidazole, pH 7.5, 25 mM KCl, 4 mM MgCl_2_, 1 mM EGTA, 1 mM DTT) incubated for 4 minutes. Excess biotin-BSA was washed out, and the surface was incubated with AB for 2 minutes. Neutravidin at 0.1 mg/mL in AB was incubated for 4 minutes. AB was added to wash out excess neutravidin. Nanotubes were added at 2–5 nM concentration in AB.nt (AB + 5–10 nM random DNA nucleotide mix) and incubated for 4 minutes. Excess nanotubes were washed out of the chamber with AB.nt. For attachment of β-cardiac myosin with eGFP, GFP nanobody SNAP at ~0.5-1 μM in AB.nt was incubated in flow chambers for 4 minutes. Excess GFP nanobody SNAP was washed out with AB.BSA.nt (AB.nt + 1 mg/mL BSA), and the surface was incubated with AB.BSA.nt for 2 minutes. For assays with cMyBP-C, N-terminal fragments at ~0.2-0.6 μM in AB.BSA.nt were incubated for 4 minutes, then washed out before the flow cell was incubated with AB.BSA.nt for 2 minutes. Finally, human β-cardiac myosin at ~0.6-0.8 μM was incubated on the surface for 4 minutes and washed out with AB.BSA.nt (for experiments with regulated thin filaments, flow cell was washed with pCa 5 buffer containing an appropriate CaCl2 concentration calculated using Max Chelator from UC-Davis). The final actin imaging solution was added (see Standard Nanotube Assay section), as described in the standard nanotube assay above. For each flow cell, the actin and nanotube channels were imaged as described above. Nanotubes with Cy3-labeled oligonucleotides incorporated into the annealing protocol and annealed to protein binding sites were imaged and used as a labeling control for Cy3-labeled cMyBP-C fragments. Nanotubes were imaged and the Cy3 and Cy5 intensities were quantified by manually selecting individual nanotubes using ImageJ and normalizing the Cy3 intensity by the corresponding Cy5 intensity for each detected pixel.

### Actin Trajectory Analysis

Actin trajectories were analyzed using the ImageJ MTrackJ plug-in. Actin movies were corrected for any drift with the TurboReg plug-in. Actin and nanotube channels were merged, and movies were analyzed to identify actin-nanotube gliding events with filaments that move along DNA nanotubes for at least 4 frames (~1 sec). For the β-cardiac myosin nanosurf assay, to clearly quantify filaments traveling on the nanotubes, we quantified velocities of the filaments that were first traveling on the coverslip surface, encountered a nanotube, and then turned sharply to glide across the nanotube. We recorded the velocity of the filament while it was on the nanotube surface as the nanotube velocity and the velocity of the filaments traveling on the surrounding coverslip surface in the nanosurf assay as nano-surface velocity.

### Statistical Analysis

Data are represented as mean values of pooled filaments ± SEM using n=82-514 filaments per condition. Experiments were independently conducted at least three times from 3-8 independent protein preparations per condition. Statistical analysis was performed using Origin 9 (OriginLab Corporation, Northampton, MA). Statistical significance was calculated for individual experiments using paired Student’s t test. Data were pooled for each condition and paired or unpaired Student’s t tests were conducted to evaluate significance. One-way ANOVA with a Tukey’s posttest was performed to assess significance when evaluating comparisons between multiple conditions (Figures 3d, 4c, and 4d) with P values *P ≤ 0.05 and ***P ≤ 0.001.

## RESULTS

### Nanosurf Motility Assay

We previously highlighted the use of synthetic nanotube thick filaments as a tool to characterize the behavior of ensembles of myosin motor proteins (34). These synthetic filaments are derived from DNA nanotubes with myosins spaced 14 or 28 nm apart. The 14 nm pattern models the spacing between consecutive actin–myosin interactions occurring in a sarcomere, where a thin filament is surrounded by three interacting thick filaments. In contrast, the spacing between myosin motors from a thick filament operating on a single actin filament is 43 nm. Our previous study primarily utilized two different highly processive unconventional myosins, V and VI. We did report an initial characterization of nanotube motility with recombinant β-cardiac myosin S1, demonstrating proof-of-concept of this assay for muscle myosins. However, the broader application of this assay to cardiac myosin was limited by the infrequent actin gliding events, consistent with the low duty ratio of cardiac myosins (Figure 1b). Therefore, in this study we developed the nanosurf assay, with myosin present on both the nanotube and the surrounding coverslip surface (Figure 1c). The presence of myosin on the coverslip surface recruits actin filaments to the surface and greatly increases the probability that a motile actin filament will encounter a nanotube. The previous format of the assay, with myosin restricted to the nanotube, exhibited only a few actin filaments per camera field-of-view (Figure 1b). In contrast, the nanosurf assay provides about a two order-of-magnitude increase in motile events along DNA nanotubes during the ~2 min data acquisition time (Figure 1c). Bi-directional motility of actin filaments is observed along DNA nanotubes, consistent with actin filaments gliding in opposing directions determined by filament polarity when interacting with nanotubes (Supplemental Video 1). To distinguish motility surface filaments from those gliding on the DNA nanotube (i.e. nanotube motility), analysis of nanotube motility events were limited to actin filaments that travel across the coverslip surface and then turn sharply to glide across the nanotube (Figure 1c schematic, *green arrow*). The thickness of actin filaments and individual DNA nanotubes (~ 10 nm) are significantly lower than the optical resolution limit of our standard epi-fluorescence microscope (~ 200 nm). Hence, our assay cannot distinguish between motile events driven by myosins on the nanotube, compared to those on the surface adjacent (~ 100 nm) but unconnected to the nanostructure. Therefore, the enhanced motility on nanotubes observed in our nanosurf assay could stem from filaments gliding on the surface myosin adjacent to the nanotube, rather than being driven by interactions with the myosins on the nanotubes themselves. To test this possibility, we compared motility on nanotubes labeled with two distinct oligos (a and b), only one of which is complementary to the oligo on the eGFP nanobody (a’) (Supplementary Figure 2a). Assays under otherwise identical conditions quantified both the number of nanotube gliding events per field of view (Supplementary Figure 2b), and the fraction of actin filament encounters with nanotubes that result in sharp turn-and-glide events (Supplementary Figure 2c). DNA nanostructures patterned with myosin (a-a’) exhibit ~ 7-fold higher motile events per field of view compared to control nanostructures patterned with a non-complementary oligo (b-a’). Hence, we conclude that ~ 90% of the nanotube gliding events stem from interactions of actin filaments with myosins patterned on the DNA nanostructure. Further, actin filaments encountering a DNA nanotube patterned with myosin (a-a’) exhibit an equal probability of either crossing over the nanotube or turn-and-glide along the nanotube (fraction of gliding events ~ 0.5; Supplementary Figure 2c). In contrast, ~ 90% of actin filaments encountering a DNA nanotube without myosin (b-a’) cross over the nanotube. Therefore, the turn-and-glide events in the control nanotubes (b-a’) are not limited by encounters with surface gliding actin filaments. Taken together, these observations demonstrate that the turn-and-glide events observed along DNA nanotubes are driven primarily by myosins patterned on the DNA nanostructure.

In the nanosurf assay, the velocity of an F-actin filament traveling along a nanotube, with an ensemble of human β-cardiac myosin HMM spaced at 14 nm intervals (0.30 ± 0.01 μm·s^−1^; n=132 pooled filaments from four independent protein preparations), is similar to the velocity of an F-actin filament traveling across the human β-cardiac myosin HMM coated coverslip surface (nano-surface; 0.31 ± 0.01 μm·s^−1^; n=139 filaments from four protein preparations). However, these velocities were significantly lower than those obtained, using matched β-cardiac myosin preparations, using a standard *in vitro* motility surface gliding assay (Figure 1d “surface”, *light grey*; 0.57 ± 0.01 μm·s^−1^; n=390 filaments). Hence, to account for any surface effects on actin gliding speeds, all future comparisons are made using matched nanosurf assays with equivalent assay conditions. These trends were also observed with fully activated (pCa 5), calcium-regulated, native thin filaments with either β-cardiac myosin HMM and or β-cardiac S1 (data from three independent protein preparations of each subfragment) (Figure 1d). Notably, ~ 2-fold higher speeds were observed with S1 compared to HMM fragments, consistent with a previously reported 2-fold higher actin-activated ATPase activity for S1 (45). With β-cardiac myosin HMM, thin filaments had an *in vitro* motility surface velocity of 0.93 ± 0.01 μm·s^−1^ (n=300 filaments), a nano-surface velocity of 0.65 ± 0.01 μm·s^−1^ (n=206 filaments), and a nanotube velocity of 0.61 ± 0.01 μm·s^−1^ (n=206 filaments). The increased velocity for thin filaments when compared to F-actin (Figure 1d), regardless of the myosin subfragment, has been described previously (46). With β-cardiac myosin S1, the regulated thin filaments had an *in vitro* motility surface velocity of 1.91 ± 0.01 μm·s^−1^ (n=359 filaments), a nano-surface velocity of 1.55 ± 0.02 μm·s^−1^ (n=270 filaments), and a nanotube velocity of 1.42 ± 0.02 μm·s^−1^ (n=270 filaments). In contrast, the average velocity of F-actin filaments in a standard *in vitro* motility surface assay with β-cardiac myosin S1 was 1.24 ± 0.01 μm·s^−1^ (n=360 filaments), once again demonstrating slower velocities associated with F-actin when compared to activated thin filaments. Taken together, these data provide baseline motility conditions for the nanosurf assay and establish the nanosurf assay as a robust method to study β-cardiac myosin ensembles.

### Motor Spacing Does Not Impact β-Cardiac HMM Motility

Next, we examined the impact of β-cardiac myosin motor spacing on actin gliding speeds. In our previous study with myosin V, myosin VI, and β-cardiac myosin S1, in a standard nanotube assay, we found no significant differences in nanotube ensemble velocity when myosin was spaced at 14 nm compared to 28 nm (34). Using the nanosurf assay with human β-cardiac myosin HMM or S1 spaced at either 14 nm or 28 nm intervals (Figure 2a,b), the velocities of either F-actin or regulated thin filaments traveling on nanotubes were quantified (Figure 2c). With HMM, we found no significant difference between F-actin velocities (data from five independent protein preparations) at 14 nm (0.30 ± 0.01 μm·s^−1^; n=162 filaments) versus 28 nm (0.32 ± 0.01 μm·s^−1^; n=143 filaments), as visually represented in the kymographs in Figure 2

**Figure 2:**
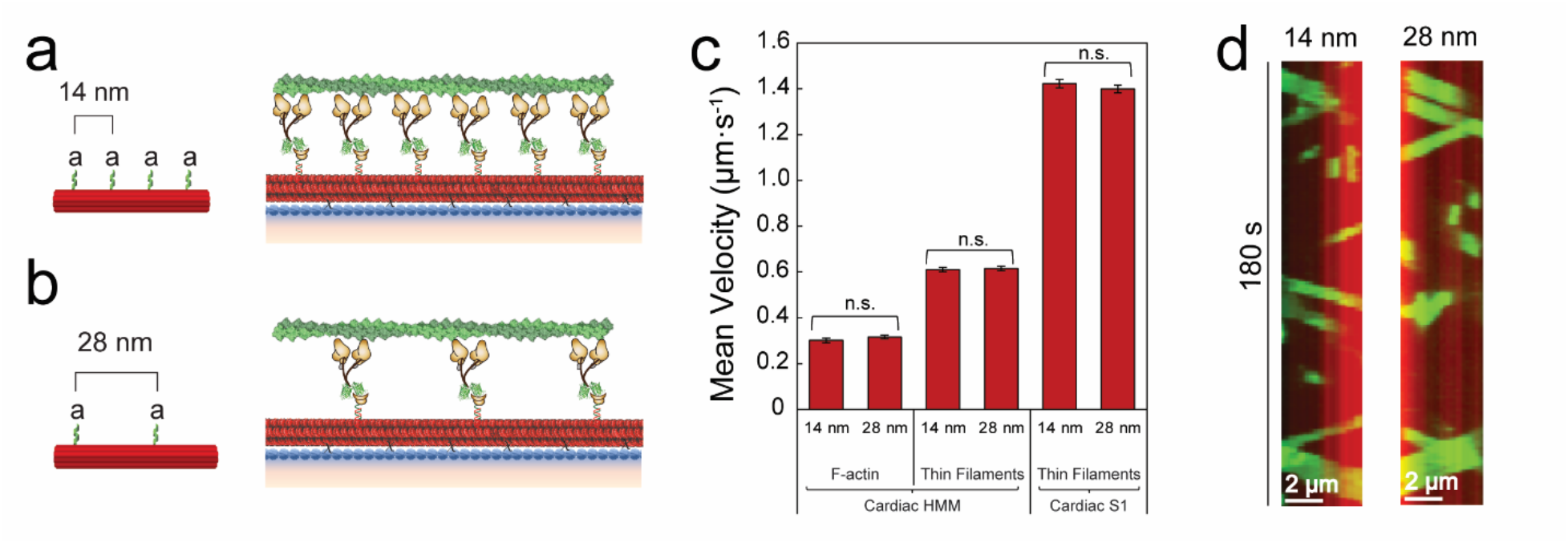
Motor Spacing Does Not Impact β-Cardiac HMM Motility. Schematic of the synthetic thick filament with nanotube motor spacing of 14 nm (**a**) or 28 nm (**b**). **c**, Velocities of F-actin (*left*) and regulated thin filaments (*middle*) on nanotubes decorated with β-cardiac myosin HMM or regulated thin filaments on nanotubes decorated with β-cardiac myosin S1 (*right*) at 14 or 28 nm spacing in the nanosurf assay. **d**, Kymographs of F-actin filaments (green) traveling on a nanotube (red) with β-cardiac myosin HMM at 14 nm (*left*) or 28 nm (*right*) spacing. Mean velocities represented as μm·s^−1^ ± SE. N = 143-270 filaments from 3-5 independent protein preparations per condition. Comparisons were performed using Student’s t-test. n.s. = not significant

Similarly, there was no significant difference in regulated thin filament velocities at pCa 5 with human β-cardiac myosin HMM or S1 (data from three independent protein preparations of each subfragment). With the HMM spaced at 14 nm versus 28 nm, regulated thin filament velocities were 0.61 ± 0.01 μm·s^−1^ (n=206 filaments) and 0.62 ± 0.01 μm·s^−1^ (n=175 filaments), respectively. With S1 spaced at 14 nm versus 28 nm, regulated thin filament velocities were 1.42 ± 0.02 μm·s^−1^ (n=270 filaments) and 1.40 ± 0.02 μm·s^−1^ (n=240 filaments), respectively. These data indicate that 28 nm spaced myosins on nanotubes can be used with interdigitated cMyBP-C so that any effects of cMyBP-C on actin filament velocities cannot be attributed to myosin density/spacing on the nanotube.

### C0-C2 Inhibits β-Cardiac HMM and S1 Nanotube Motility

We used the nanosurf assay as a tool to investigate the role of cMyBP-C interactions with actin and myosin, in regulating muscle contraction. The cMyBP-C protein contains 11 domains, C0-C10, with the N-terminal domains C0-C2 thought to be most critical for its regulatory interactions with actin and myosin (Figure 3a) (10, 17, 18). Further, the M-domain linker between the C1 and C2 domains contains 4 phosphorylatable serines that, once phosphorylated in response to adrenergic stimuli, may modulate cMyBP-C’s impact on cardiac contractility (28–32). Hence, the C0-C2 N-terminal fragment was used to simulate the influence of the full-length cMyBP-C. Human β-cardiac myosin spaced at 28 nm intervals (Figure 3b, *myosin only*) on the nanotube was interdigitated with C0-C2, also at 28 nm intervals, (Figure 3b, *+C0-C2*) containing either no ER/K α-helical linker, a 10 nm ER/K α-helical linker, or a longer 30 nm ER/K α-helical linker to potentially facilitate cMyBP-C interactions with both actin and myosin (Supplemental Figure 1b,c). The surface around the nanotubes was blocked with BSA prior to the introduction of C0-C2 (see Methods). Hence, C0-C2 fragments should bind primarily to the nanotubes. However, a ~ 30% reduction in surface actin gliding velocities were observed, consistent with residual binding to the nano-surface (Supplemental Figure 3). Nonetheless, when C0-C2 with a 30 nm ER/K linker was present on a nanotube decorated with β-cardiac myosin HMM (data from eight independent protein preparations), ensemble velocity of F-actin was reduced 4-fold from 0.29 ± 0.01 μm·s^−1^ (n=211 filaments) to 0.07 ± 0.002 μm·s^−1^ (n=389 filaments) (Figure 3c; Supplemental Video 2). Likewise, with the presence of C0-C2, regulated thin filament velocity at pCa 5 on β-cardiac myosin HMM nanotubes (data from three independent protein preparations) was reduced 6-fold from 0.60 ± 0.02 μm·s^−1^ (n=171 filaments) to 0.10 ± 0.003 μm·s^−1^ (n=223 filaments). These effects are comparable to a previously reported ~ 5-fold decrease in actin gliding speeds in a standard *in vitro* motility assay with chicken skeletal muscle myosin and cMyBP-C adsorbed at high concentrations (0.4 μM) (30).

**Figure 3:**
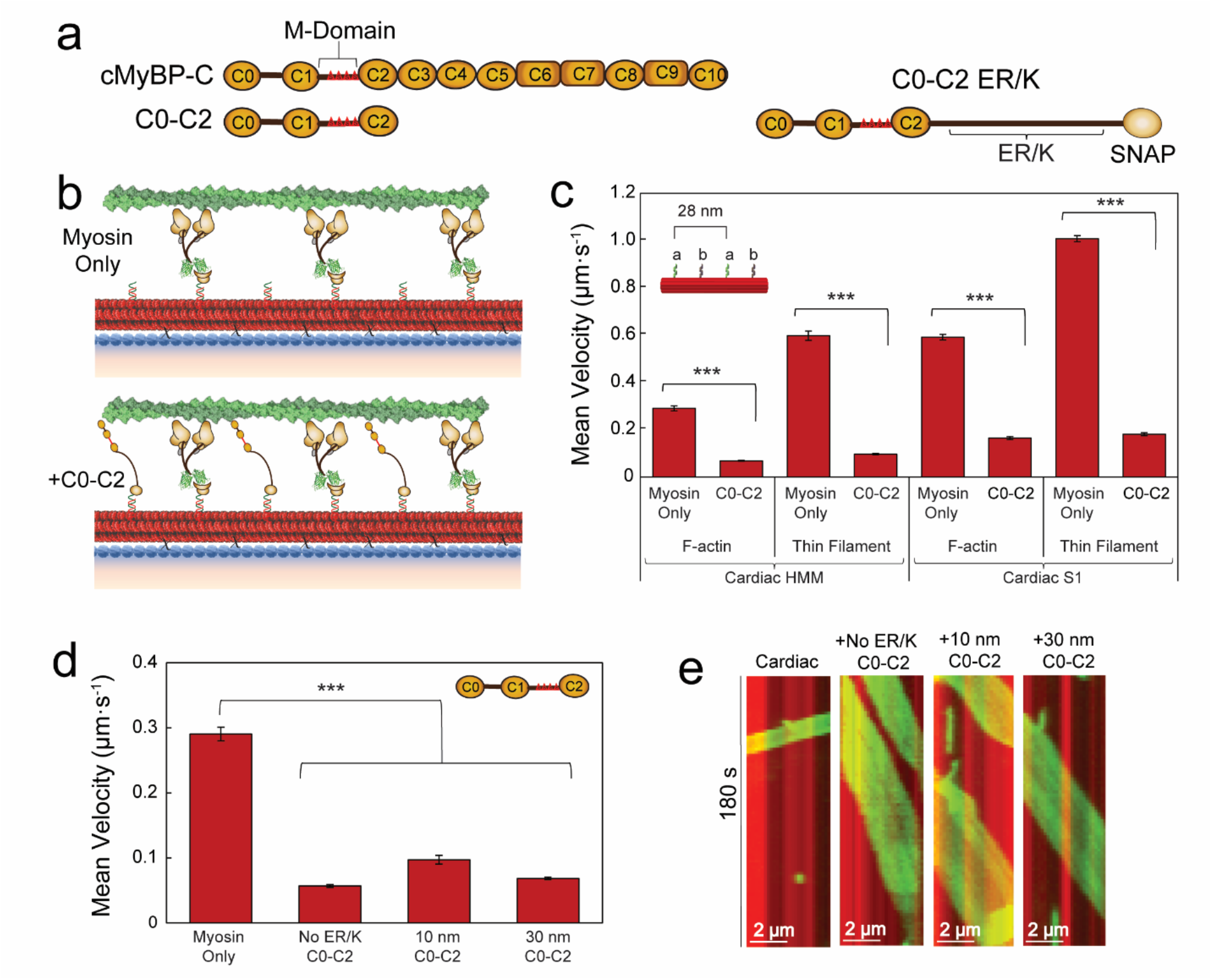
C0-C2 Inhibits β-Cardiac HMM and S1 Nanotube Motility. **a**, Schematic of cMyBP-C domains C0-C10 with the M-domain (*red triangles*) containing 4 phosphorylatable serines in the linker region between the C1 and C2 domains and the N-terminal fragment used, C0-C2. C0-C2 schematic (*right*) shown with the encoded ER/K linker and SNAP tag. **b**, Schematics of synthetic thick filaments with β-cardiac myosin HMM bound to oligo a’ at 28 nm intervals (myosin only, *top*) and interdigitated C0-C2 containing M-domain, bound to oligo b’ (*bottom*). **c**, Velocities of F-actin and regulated thin filaments on nanotubes decorated with β-cardiac myosin HMM (*left*) or β-cardiac myosin S1 (*right*) bound to oligo a’ alone versus myosin +C0-C2 containing a 30 nm ER/K bound to oligo b’ in the pattern shown (*inset*). **d**, Mean velocities of F-actin filaments on nanotubes decorated with β-cardiac myosin HMM only versus β-cardiac HMM nanotubes with interdigitated C0-C2 (*inset*) containing no ER/K helix, a 10 nm ER/K helix, or a 30 nm ER/K helix. **e**, Kymographs of F-actin filaments (*green*) traveling on a nanotube (*red*) with β-cardiac myosin HMM only versus cardiac nanotubes with interdigitating C0-C2 containing no ER/K helix, a 10 nm ER/K helix, or a 30 nm ER/K helix. Mean velocities represented as μm·s^−1^ ± SE. N = 82-514 filaments from 3-8 independent protein preparations per condition. Significance calculated using Student’s t-test or one-way ANOVA where appropriate. Significance is denoted as \*\*\**P* ≤ 0.001

We found no significant impact of the ER/K linker or linker length on the inhibitory capacity of C0-C2, as all C0-C2 ER/K constructs inhibited F-actin nanotube motility ~4-fold (Figure 3d; data from n=82-389 filaments pooled from 3-8 independent protein preparations), as visually represented in the kymographs in Figure 3e. Together, these studies demonstrate the robust use of the nanosurf assay to recapitulate the effects of cMyBP-C on actomyosin motility.

To distinguish between cMyBP-C effects being imparted through its interaction with actin and/or myosin, we utilized nanotubes decorated with β-cardiac myosin S1, which lacks the S2 domain that has been implicated as the primary cMyBP-C interacting region in myosin (19–23). β-cardiac S1 and HMM displayed similar velocity inhibition with the C0-C2 fragment (Figure 3c). Specifically, the presence of C0-C2 with β-cardiac S1 (data from three independent protein preparations) reduced F-actin velocity almost 4-fold (0.59 ± 0.01 μm·s^−1^ to 0.16 ± 0.01 μm·s^−1^; n=180 and 267, respectively) and regulated thin filament velocity almost 6-fold (1.01 ± 0.01 μm·s^− 1^ to 0.18 ± 0.01 μm·s^−1^; n=450 and 514, respectively). These data suggest that C0-C2 interactions with actin, rather than myosin S2, are the primary determinant of its effects on actin gliding speed in the nanosurf assay. The significance of C0-C2-actin interactions is also evident in the extent of actin recruitment to nanotubes. Unlabeled nanotubes do not recruit actin filaments (Supplementary Figure 4a). When the C0-C2 fragment was attached to the nanotubes without motor, it pulled more F-actin filaments (237.0 filaments per field of view ± 37 standard deviation; Supplemental Figure 4b) onto nanotubes than motor in the absence of C0-C2 (146.7 filaments per field of view ± 22 standard deviation; Supplemental Figure 4c). The presence of both C0-C2 and β-cardiac myosin HMM on nanotubes yielded greater actin recruitment on nanotubes than either of them alone (326.7 filaments per field of view ± 116 standard deviation; Supplemental Figure 4d). Robust actin recruitment to nanotubes was still seen by C0-C2, even when the surface was blocked in the standard nanotube assay (Supplemental Figure 4e).

### cMyBP-C M-domain Phosphorylation and Structurally Dependent Inhibition of β-Cardiac Myosin HMM and S1 Nanotube Motility

To investigate the role of individual cMyBP-C domains in the inhibition of actomyosin motility, we examined alternative N-terminal fragments, C0-C1f and C1-C2 in our nanosurf assay (Figure 4a). The C0-C1f fragment contains the C0 and C1 domains, as well as the first 17 amino acids of the M-domain (Figure 4a). This segment of the M-domain contains several arginine residues proposed to be essential for cMyBP-C interaction(s) resulting in actomyosin inhibition (Figure 4b, *left, inset*) (30, 47). However, it lacks the four phosphorylatable serines found in the M-domain (Figure 4a) (30, 48). The C1-C2 fragment contains the C1 and C2 domains, as well as the entire M-domain linker region between C1 and C2. Using the nanosurf assay, human β-cardiac myosin spaced at 28 nm intervals on the nanotube was interdigitated with either C0-C1f (Figure 4b, *left*), C1-C2 (Figure 4b, *right*), or C0-C2, each spaced at 28 nm. The presence of C0-C1f did not significantly impact F-actin or thin filament motility driven by either β-cardiac myosin HMM or S1 (Figure 4c; data from n=100-274 filaments pooled from 3-4 independent protein preparations). In contrast to C0-C1f, the C1-C2 fragment was as inhibitory of F-actin and thin filament motility as C0-C2 (Figure 4d). Specifically, both C1-C2 and C0-C2 reduced F-actin velocity on β-cardiac myosin HMM nanotubes ~ 4-fold, and reduced thin filament velocity on β-cardiac myosin S1 nanotubes ~8-fold (Figure 4d, e; data from n=90-270 filaments pooled from 3-4 independent protein preparations). These data are consistent with similar inhibitory effects observed with the addition of C1-C2 or C0-C2 to a standard *in vitro* motility assay (30, 33).

**Figure 4:**
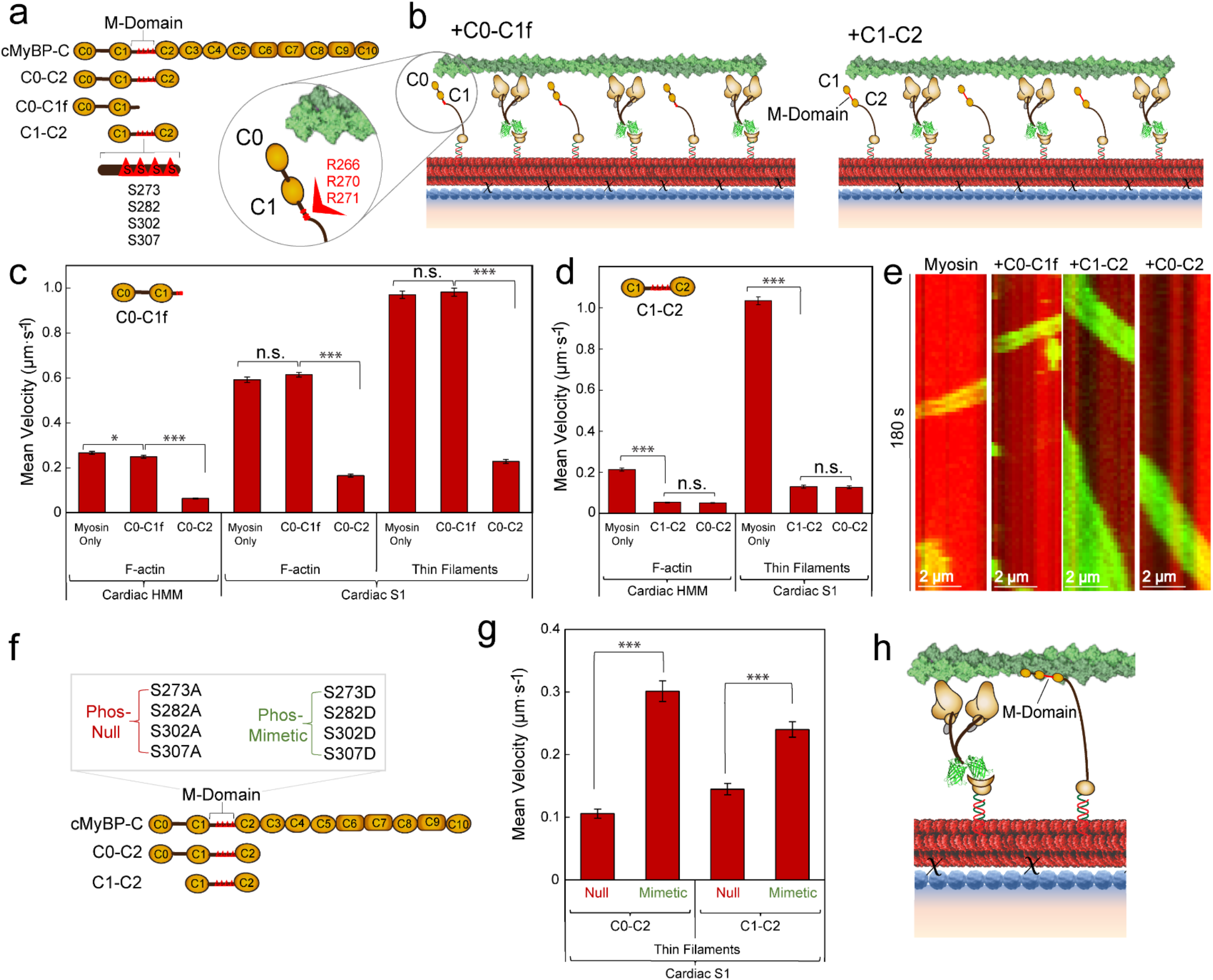
cMyBP-C M-domain Essential for Inhibition of β-Cardiac Myosin HMM and S1 Nanotube Motility. **a**, Schematic of cMyBP-C domains C0-C10 containing the M-domain (*red triangles*) in the linker region between the C1 and C2 domains and the N-terminal fragments used, including C0-C2, C0-C1f, and C1-C2. The four phosphorylatable serines in the M-domain are represented by red triangles (S273, S282, S302, S307). **b**, Schematics of synthetic thick filaments with β-cardiac myosin HMM bound to oligo a’ at 28 nm intervals and interdigitated C0-C1f (*left*) or C1-C2 (*right*) containing the entire M-domain, bound to oligo b’. C0-C1f (*inset*) contains the first 17 amino acids of the M-domain including several arginine residues (R266, R270, R271) oriented away from the actin filament. **c**, Velocities of F-actin on nanotubes decorated with β-cardiac myosin HMM (*left*) and either F-actin (*middle*) or regulated thin filaments (*right*) on nanotubes decorated with β-cardiac myosin S1. β-cardiac nanotubes were either labeled with the motor alone or interdigitated with C0-C1f or C0-C2 containing 30 nm ER/K helices. **d**, Velocities of F-actin on nanotubes decorated with β-cardiac myosin HMM (*left*) and regulated thin filaments on nanotubes decorated with β-cardiac myosin S1 (*right*) versus β-cardiac nanotubes with interdigitated C1-C2 or C0-C2 containing 30 nm ER/K helices. **e**, Kymographs of F-actin filaments (*green*) traveling on a nanotube (*red*) with β-cardiac myosin HMM alone versus cardiac nanotubes with interdigitating C0-C1f, C1-C2, or C0-C2. **f**, Schematic of cMyBP-C domains C0-C10 with the M-domain (*red triangles*) containing 4 phosphorylatable serines in the linker region between the C1 and C2 domains mutated to alanines (phospho-null) or aspartic acids (phospho-mimetic) within the C0-C2 and C1-C2 N-terminal fragments. **g**, Mean nanotube velocities of regulated thin filaments on nanotubes decorated with β-cardiac myosin S1 and interdigitating phospho-null or phospho-mimetic C0-C2 (*left*) or C1-C2 (*right*) containing 30 nm ER/K helices. **h**, Schematic of β-cardiac myosin (brown) synthetic thick filament with cMyBP-C N-terminal fragment (yellow) bound to actin (green), functioning as a tether between thick and thin filaments and reducing velocity. Mean velocities represented as μm·s^−1^ ± SE. N = 90-410 filaments from 3-4 independent protein preparations per condition. Significance was calculated using Student’s t-test. Significance is denoted as \**P* ≤ 0.05, \*\*\**P* ≤ 0.001.

The C1-C2 fragment contains the entire M-domain, including the phosphorylatable serines (S273, S282, S302, S307) that regulate cardiac contractility. Hence, we generated phospho-null (Ser to Ala) and phospho-mimetic (Ser to Asp) C0-C2 and C1-C2 N-terminal fragments (Figure 4f), as previously reported for the C0-C3 fragment (1, 48). Mass spectrometry suggests that our insect-cell expressed N-terminal fragments are not phosphorylated (*data not shown*). The phospho-null C0-C2 and C1-C2 fragments inhibited thin filament velocity over β-cardiac myosin S1 nanotubes to a similar extent as unmodified fragments (Figure 4g; data from n=240-270 filaments pooled from three independent protein preparations). However, the phospho-mimetic C0-C2 and C1-C2 mutants exhibited less inhibition (Figure 4g), as previously reported for the identical phosphomimetic substitutions in the C0-C3 fragment used in the *in vitro* motility assay (1, 48). These results highlight the importance of the phospho-serines in regulating cMyBP-C mediated inhibition of β-cardiac myosin function.

## DISCUSSION

In this study, we utilized DNA nanotechnology to create a synthetic myosin thick filament with incorporated cMyBP-C. This synthetic thick filament controls the stoichiometry and spatial configuration of protein-protein interactions to provide mechanistic insights into cMyBP-C function. Using synthetic thick filaments with an assortment of cMyBP-C N-terminal fragments, we characterized the effects of cMyBP-C domains on velocity of either F-actin or regulated thin filaments. Our findings suggest that the binding of cMyBP-C to actin may underly the inhibitory effect of cMyBP-C on actomyosin motility over nanotubes (Figure 4h) and that the cMyBP-C M-domain is crucial to this inhibition. Further, phosphomimetic cMyBP-C fragments, displaying diminished inhibition of thin filament velocity on nanotubes, reproduce the established modulatory role of M-domain phosphorylation on cMyBP-C function (1, 28, 29, 49). Hence, while the synthetic thick filament does not capture the inherent three-dimensional nature of the cardiac muscle sarcomere and the strain-dependent activation of myosin (50), our study demonstrates a robust approach to investigate subtle changes in contractility through genetically inherited mutations and/or physiological regulation through post-translational modifications.

*In vitro* motility assays with native thick filaments from mouse cardiac tissue maintain the myosin to cMyBP-C stoichiometry and spatial relationships. These were critical to establish that cMyBP-C mediates thin filament slowing and calcium sensitization within the C-zone. However, such native thick filament assays are not suitable for molecular structure-function studies with systematic perturbations to the myosin or cMyBP-C, due to the high cost of animal model design. Our nanosurf assay fills this need by defining the stoichiometry and spatial relations between myosin and cMyBP-C and the relative ease in expression of human β-cardiac myosin and cMyBP-C fragments that decorate the nanotube surface.

The nanosurf assay also overcomes a key limitation of our previously reported standard nanotube assay (34). The standard nanotube assay is well suited to study high duty ratio motors, which readily recruit actin filaments from solution and display abundant motility. In contrast, the use of low duty ratio, β-cardiac myosin nanotubes is limited by infrequent engagement of actin filaments from the surrounding solution (Figure 1b). To overcome this challenge, we seeded β-cardiac myosin subfragments at a sufficient density on the surface to which the nanotubes were attached (see Methods) so that the coverslip surface myosin recruited actin filaments from solution, which then encountered the neighboring nanotubes, substantially enhancing the frequency of nanotube motile events (Figure 1c). The actin gliding speeds on nanotubes and the surrounding surface are indistinguishable, however, these speeds are significantly reduced compared to that observed in the standard *in vitro* motility assay (Figure 1d). These data suggest that the surface conditions and/or the mode of myosin attachment likely contribute to the lower speeds in the nanosurf assay. Specifically, the myosin attachment strategy (see Methods) may reduce the efficient transfer of myosin displacements to the actin filament, leading to slower velocities. Hence, we focused on motility comparisons between conditions only within the nanosurf assay. In addition, similar flexibility by the addition of a variable length ER/K α-helical linker to the cMyBP-C fragment ensured that cMyBP-C on the nanotube was spatially free to interact with β-cardiac myosin and/or actin filaments (Supplemental Figure 2b,c). However, the ER/K α-helical linker was not essential for the observed cMyBP-C effects on actin filament motility, as constructs lacking the ER/K α-helical linker demonstrated equivalent inhibition. Overall, the multiple rotational elements in the myosin and cMyBP-C attachment, combined with myosin’s access to numerous sites along the actin filament (Supplemental Figure 2c), yields significant conformational freedom without limiting protein-protein interactions.

We observed that slowing of F-actin or fully activated thin filament velocity by the C0-C2 N-terminal fragment was structurally localized to the C1-C2 domains, with the M-domain linker between C1 and C2 being a critical component (Figure 4d). The importance of the M-domain is emphasized by phosphomimetic replacement of four serines within the M-domain that reduce the observed slowing of thin filament velocity, as reported previously (1, 48). To further characterize this regulatory domain, we utilized the C0-C1f fragment, which contains only the first 17 amino acids of the M-domain but lacks the four phosphorylatable serines (30, 48). Interestingly, a previous study showed inhibition of thin filament velocity by C0-C1f in a standard *in vitro* motility assay, and it was proposed that several C-terminal arginine residues in this fragment may be particularly important for this inhibition (Figure 4b, *inset*) (30, 47). The inability of the C0-C1f to slow thin filament velocity in our nanosurf assay (Figure 4c), despite the presence of several C-terminal arginine residues (Figure 4b, *inset*), further supports the importance of the entire M-domain. A potential source for this discrepancy is the weaker binding of C0-C1f to actin that manifests only when high concentrations of this fragment are used in the standard *in vitro* motility assay (30).

*In vitro*, both actin and myosin-binding have been described for the cMyBP-C N-terminus (17–19, 21, 22, 24–27). Whether one or both of these binding partner interactions is responsible for the modulatory capacity of cMyBP-C is still a matter of debate. The myosin S2 domain was the first structural element identified in cMyBP-C binding (19). Using the β-cardiac S1 fragment, which is devoid of the S2 segment, on the nanotubes, substantial slowing of actin gliding speeds in the presence of the C0-C2 and C1-C2 fragments remained (Figure 4d). Although we cannot rule out binding of these fragments to the myosin regulatory light chain (22) or to a yet defined region of the motor domain (32), the preponderance of data both here and in the literature (1, 9, 17, 18, 24–29) suggest that the cMyBP-C inhibitory effect *in vitro* may in large part be due to its interaction with actin. In fact, recent *in vivo* super-resolution data suggest that the cMyBP-C N-terminus in mouse myocardium resides predominantly near the thin filament regardless of the muscle’s active state and that only when activated would the myosin head domain being in close enough proximity to interact with the cMyBP-C terminus (18). The slowing of thin filament velocity may arise, in part, due to drag forces imposed by cMyBP-C binding and tethering the thin filament to the nanotube (51), thereby slowing actomyosin kinetics (52, 53). In addition, cMyBP-C may be competing with myosin for binding sites on actin, thus reducing the number of force generating motors, as suggested previously (25, 54–56).

## CONCLUSION

In conclusion, we have established the DNA synthetic thick filament nanosurf assay as a robust tool for the characterization of β-cardiac myosin and cMyBP-C interactions. We have recapitulated actomyosin inhibition by cMyBP-C N-terminal fragments on recombinant human β-cardiac myosin DNA nanotubes and begun to dissect the mechanisms underlying contractile regulation by cMyBP-C. Our results emphasize the importance of the M-domain and its phosphorylation in regulating actomyosin motility. Our study also highlights the role of cMyBP-C N-terminal interactions with actin in regulating motility and suggests the actin-cMyBP-C interaction is the dominant interaction underlying actomyosin inhibition by cMyBP-C. Taken together, our results support a model where cMyBP-C tethers the thick and thin filaments at high calcium, imposing drag, and slowing thin filament velocity. Future studies can build on this foundation to examine other aspects of contractile regulation, including external load, on cMyBP-C function.

## AUTHOR CONTRIBUTIONS

A.M.T., A.R., D.M.W., C.M.Y., and S.S. designed research. A.M.T., W.T., D.V.R., D.V., A.R., and S.B.P. performed research. C.M.Y. and D.M.W. provided key reagents. A.M.T., D.V., A.R., D.M.W., C.M.Y., and S.S. analyzed data. The manuscript was written by A.M.T., D.M.W., C.M.Y., and S.S. and approved by all authors.

## ACKNOWLEDGMENTS

The authors would like to thank Ruth Sommese for technical expertise and Michael Ritt for helpful discussions and technical assistance. Research was supported by the NIH (1R35GM126940 to S.S.; 1R01HL150953 to D.W., C.Y., and S.S.; 1F30HL146089-01 to A.T.)

