## Supplemental Files for "Dissecting β-Cardiac Myosin and Cardiac Myosin-Binding Protein C Interactions using a Nanosurf Assay"

### SUPPLEMENTAL INFORMATION

#### FIGURES:

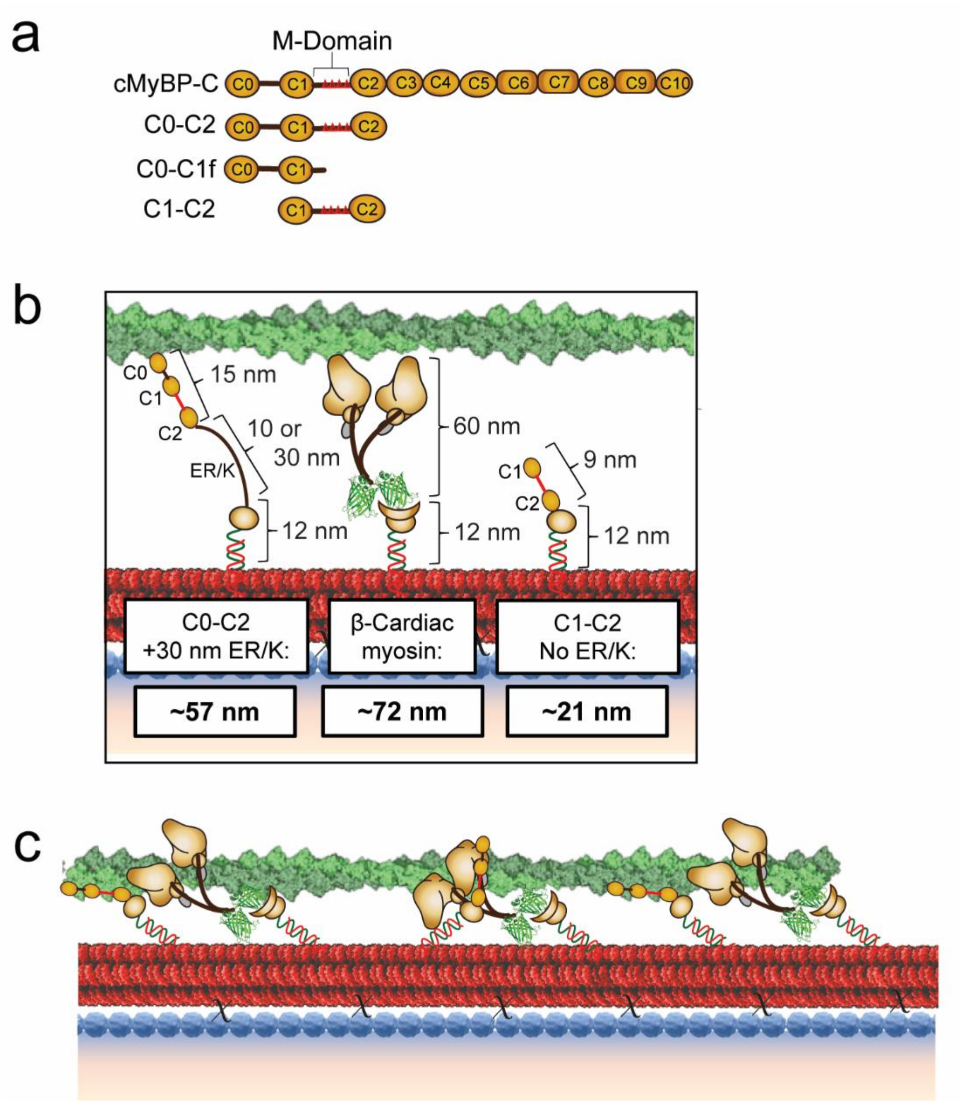


**Figure S1: Flexibility in the Nanosurf Assay allows cMyBP-C Interactions with Actin and/or Myosin.**
**a,** Schematic of cMyBP-C domains C0-C10 containing the M-domain in the linker region between the C1 and C2 domains and the N-terminal fragments used, including C0-C2, C0-C1f, and C1-C2. **b**, Diagram showing lengths of representative cMyBP-C N-terminal fragments, C0-C2 (left, *yellow;* ~15 nm) and C1-C2 (right, *yellow;* ~9 nm) and recombinant human β-Cardiac myosin HMM (center, *brown;* ~60 nm) with C-terminal GFP attached to the nanotube (*red*) via GFP nanobody-SNAP. All proteins are attached to the nanotube via a SNAP protein labeled with oligo (~12 nm) complementary to the nanotube DNA handle. C0-C2 is shown with an encoded ER/K linker (left; 10 or 30 nm). Total approximate lengths of bound proteins are listed: C0-C2 + 30 nm ER/K (~57 nm), β-Cardiac myosin (~72 nm), and C1-C2 without an ER/K (~21 nm). **c**, Diagram depicting spatially staggered interaction sites on the actin filament (*green*) and possible cMyBP-C interactions with both actin (left, right C0-C2 fragments) and myosin S2 (center C0-C2 fragment).

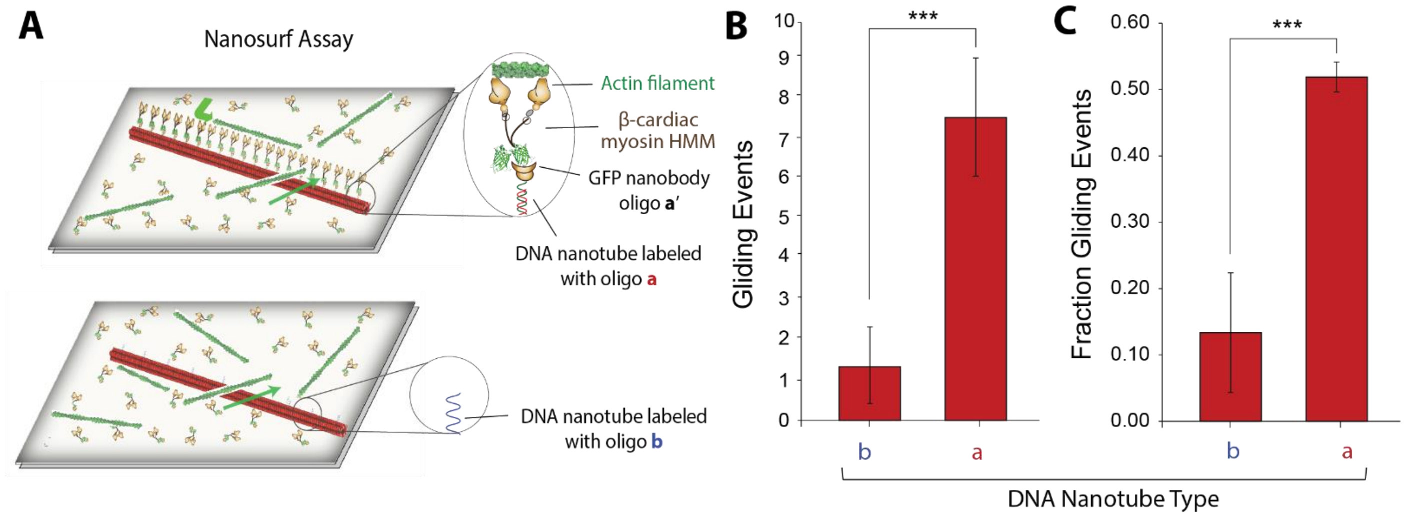


**Figure S2: Motility on DNA nanotubes in nanosurf assay is driven by myosins linked to the nanotube.** **a,** Schematic of nanosurf assay with DNA nanotubes labeled with either a or b-type oligos. GFP nanobody is labeled with an oligo a', complementary to oligo a. **b,c** Actin filaments glide on surface-anchored myosin, and either cross or turn sharply and glide along DNA nanotubes. **b,** Number of turn-and-glide events on DNA nanotubes (per field of view in 2 min video). **c,** Fraction of encounters of surface gliding actin filaments that turn-and-glide along DNA nanotubes (turn-and-glide events / (turn-and-glide + cross events)). Data are mean+/-SD of at least three movies, derived from three independent flow chambers.

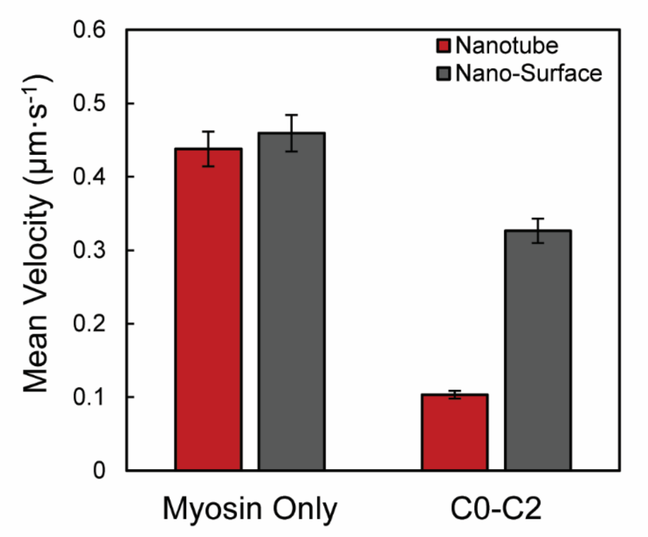


**Figure S3: C0-C2 Impact on β-Cardiac HMM Nanotube and Nano-Surface Motility.**
Velocities of F-actin on the coverslip surface in the nanosurf assay (Nano-Surface; *dark grey*) and on the nanotubes in the nanosurf assay (*red*) for nanotubes decorated with β-cardiac myosin HMM bound to oligo a' alone (*left*) versus myosin +C0-C2 containing a 30 nm ER/K bound to oligo b' (*right*). Mean velocities represented as $\mu m \cdot s^{-1}$ ± SE. N = 63-82 filaments from 3 independent protein preparations per condition.

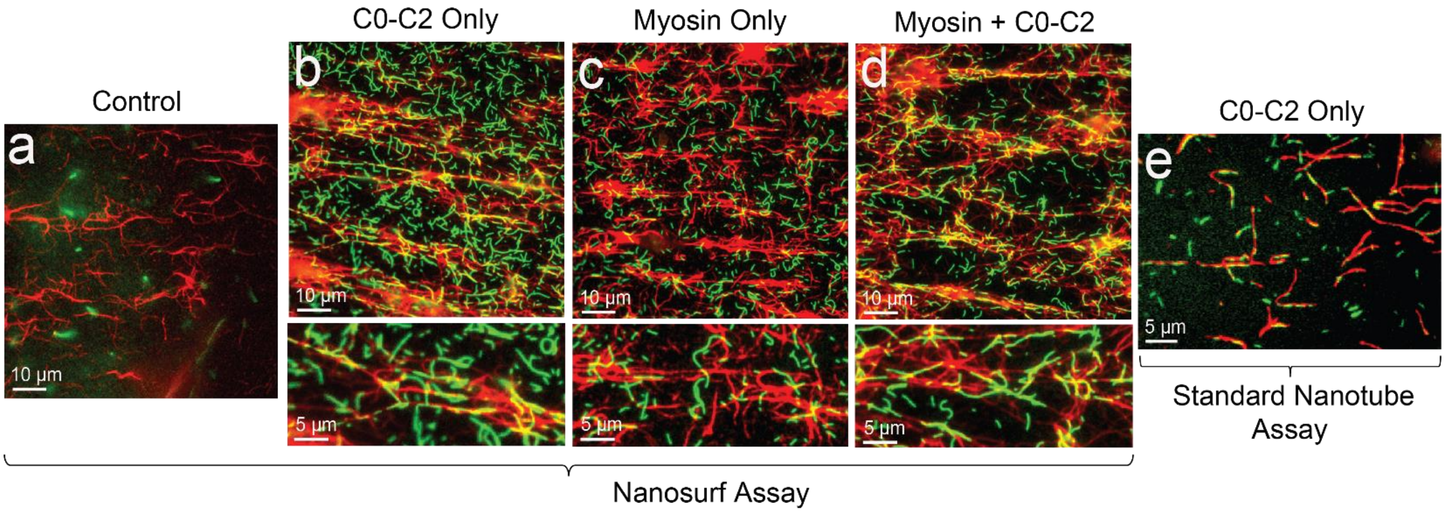


**Figure S4: cMyBP-C C0-C2 N-terminal Fragment Recruits Actin onto Nanotubes.**
**a-d,** We examined actin (*green*) recruitment onto nanotubes (*red*) in the nanosurf assay. Nanotubes were **a,** unlabelled, or labeled with **b,** C0-C2 only (containing 30 nm ER/K, with fragments spaced at 28 nm intervals), **c,** β-cardiac myosin HMM only (myosin spaced at 28 nm intervals), or **d,** β-cardiac myosin HMM + C0-C2 (C0-C2 contained 30 nm ER/K; myosin was spaced at 28 nm intervals and interdigitated with C0-C2 for a final spacing of 14 nm between myosin and C0-C2 proteins). The top panels in **a-d** show the field of view at 1000x with selected enlargements for **b-d** shown in the bottom panels. **e**, Actin recruitment was also examined using a standard nanotube assay blocking conditions and nanotubes labeled with C0-C2 + 30 nm ER/K.

**VIDEOS:**

**Video S1: Bi-direction movement of actin filaments on Nanotubes.** Representative video depicting a nanosurf assay with F-actin filaments (*green*) traveling on nanotubes (*red*) labeled with β-cardiac myosin HMM spaced 14 nm apart. Two actin filaments are seen traveling in opposite directions on the nanotube in the center.

**Video S2: Inhibition of Actin Velocity by C0-C2 bound to β-cardiac myosin HMM Nanotubes.** Representative video depicting a nanosurf assay with F-actin filaments (*green*) traveling on nanotubes (*red*) labeled with interdigitated β-cardiac myosin HMM and C0-C2 (+10 nm ER/K) spaced 14 nm apart. Actin filaments can be seen moving slower on the C0-C2-decorated β-cardiac HMM nanotubes (nanotube velocity) before accelerating onto the surrounding motility surface (nano-surface) coated with β-cardiac myosin HMM.
